# Optimizing forest canopy structure retrieval from smartphone-based hemispherical photography

**DOI:** 10.1101/2021.03.17.435793

**Authors:** Gastón Mauro Díaz

## Abstract

1. Hemispherical photography (HP) is a long-standing tool for forest canopy characterization. Currently, there are low-cost fisheye lenses to convert smartphones into highly portable HP equipment (smartphone-based HP, hereafter SHP). However, there is an obstacle to having a close-to-ideal method for citizen science and large-scale or opportunistic sampling: the known sensitivity of HP to illumination conditions. The purpose of this paper is to test a ready-to-use approach based on previous research, and to contribute to quantifying the errors associated with choosing SHP in non-recommended light conditions over well-established HP practices.
2. In 30 locations distributed in broadleaf and coniferous woodlands, a total of 1080 photographs were taken with two smartphone models, manipulating the exposure, and under varied sky conditions. After image binarization, accurate reference data was employed to evaluate the reliability of extracting canopy parameters from SHP.
3. The proposed methodology can reliably quantify canopy openness (RMSE ~ 0.04) and plant area index (RMSE ~ 10%), suggesting that SHP, when used following the recommendations from the present study, allows the retrieval of unbiased canopy metrics independently of sky conditions and forest type.

## Introduction

Ecological oriented research often characterizes vegetation canopies through the leaf area index (LAI), defined as one-half the total photosynthetic area per unit of horizontal ground surface area (Chen & Black, 1992). There is no inherently superior method for in-situ measuring of LAI, each has its own advantages and disadvantages. Currently, choosing a method to maximize accuracy is relatively straightforward, but little is known about solutions suited for projects seeking for flexibility.

Current most popular indirect methods are conceptually based on the point quadrat method, which consists of piercing the canopy in a constant direction with a long needle while registering the number of contacts between the needle point and the canopy (Nilson, 1971). A variant of this method consists of registering whether the point contacts the canopy or not. If this is repeated several times, the probability of no contact (P_0_) can be calculated. In the ideal canopy—made of ideal leaves randomly located and forming an infinite layer with their long axis randomly deviated from the north and inclined following a theoretical leaf angle distribution (LAD)—P_0_ obey a Poisson distribution parameterized by LAI, LAD, and needle inclination to the vertical (the zenith angle, *θ*). In a real canopy, P_0_ will depend on many factors, but when pretending that the observed values came from the ideal canopy, the LAI obtained by inversion is known as the effective LAI (eLAI). Metrics more sophisticated than P_0_ can be used to calculate the clumping index *(Ω*), which applied to eLAI corrects deviations from the assumption of randomness in location. The corrected index is known as the plant area index (PAI), which can be defined the same as LAI, but without distinguishing photosynthetic from non-photosynthetic area (Yan et al., 2019).

Modern methods replace the needle-like probe with virtual probes, namely light beams or lines of sight. Light beams are used in both active methods (laser beams) and passive methods (sunlight) (Yan et al., 2019). These methods require cumbersome and expensive instruments, making them unattractive for projects seeking flexibility, which rather leans towards the line-of-sight approach since it only requires an imaging device. The challenge of this approach is using the image to evaluate whether an imaginary probe starting on lens location would contact or not the canopy. Therefore, the image has to be classified into gap and non-gap. Afterward, if 1 is assigned to gap pixels and 0 to non-gap pixels (binarization), their average value (for any sky region) is known as the gap fraction (GF) and is equivalent to P_0_.

Using the line of sight as a virtual probe assumes the lens projection is known, the image is a trusty representation of the canopy (image fidelity), and it can be binarized accurately. Any violation of these assumptions is an error source. For instance, improper lens calibration, images showing missing or distorted canopy elements, and low separability between gaps and non-gaps.

There are two canopy photography methods that can be conducted with consumer-grade equipment, hemispherical photography (HP) and restricted view photography (RVP) (Yan et al., 2019). The concrete advantage of RVP over HP is the type of lens needed because the default lenses of cameras and mobile devices are not enough to conduct HP; auxiliary fisheye lenses are required to convert them into capable HP equipment—an exception is the Arietta (2021) method. Nevertheless, although RVP is experiencing a fast development, HP is a more established method. The scope of this study was upward-looking smartphone-based HP applied to fully developed canopies, but conclusions can be translated to HP in general as well as RVP.

The most tempting acquisition protocol is using the device in automatic mode under any canopy and light condition (point-and-shoot mode). However, obtaining good results with those kinds of photographs is challenging (Díaz et al., 2021). The older approach has been to do fieldwork in diffuse light conditions (overcast and twilight) to ensure data separability along a single feature (blue channel digital number), and apply a manual global threshold to binarize the image. In pursuit of objectivity and replicability, this manual approach was gradually replaced by general-purpose algorithms for classifying bimodal histograms (Jonckheere et al., 2005), but these algorithms showed a strong dependency to the exposure setting since it controls the histogram shape; therefore, the subjectivity passed from threshold determination to exposure setting, so research efforts were directed towards exposure protocols. All effective exposure protocols aim to make the sky as bright as possible without degrading the data. The most reliable protocol uses an open-sky exposure reading to correctly expose the sky even under dense canopies (Zhang et al., 2005). Because this approach is impractical, it is often replaced by histogram inspection in the field and lab, which is facilitated by RAW acquisition (Macfarlane et al., 2014). In general, the goal of exposure protocols is to produce a blue channel with an easy-to-binarize bimodal histogram.

Under perfect diffuse light conditions, the standard practice is considering pixels from the darker and brighter mountain-like curves in the histogram as plant and sky pixels, respectively, and pixels from the valley-like curves as mixed pixels. This assumes a subject made of homogeneously dark objects against a homogeneously light background. This kind of subject, known as artificial targets, has been manufactured to study algorithms performance under ideal conditions (Macfarlane et al., 2014). However, their use for studying HP in non-diffuse light conditions is limited because canopy parts reached by direct light tend to be closer in brightness to the darkest parts of the sky. This phenomenon can be observed even in RVP—although considered less sensitive to illumination conditions than HP (Yan et al., 2019). Hwang et al. (2016) reported problems to binarize photographs showing sunlit leaves, Chianucci et al. (2021) unimodal histograms that were successfully classified only after applying a color enhancement preprocessing (Díaz & Lencinas, 2015), and Lusk (2022) difficulties classifying sunlit bark and the deep blue of broken cloud skies.

Smartphones’ high-portability and widespread use puts HP sensitivity to illumination conditions as the main obstacle for having a close-to-ideal method—smartphone-based HP (SHP)—for citizen science and large-scale or opportunistic sampling. A deep learning approach has proven effective in making HP independent of illumination conditions, but it requires a laborious calibration along the line of what allometric equations require (Díaz et al., 2021). Automatic out-of-the-box solutions—like the ones applied to diffuse light acquisitions but specially designed for non-diffuse light—follow the general strategy of calculating a new feature from the RGB channels to exploit the color information domain, and then applying an automatic thresholding algorithm to this new feature (Díaz & Lencinas, 2015; Loffredo et al., 2016). The performance of this type of algorithm in different settings generated by varied light conditions, canopy structure, and camera settings has not been evaluated yet. The purpose of this paper is to evaluate the application of Díaz & Lencinas (2015) method to SHP—the comparison of this method against Loffredo et at. (2016) was done in Díaz & Lencinas (2021). The experiment involved two smartphones models, several exposure settings, two image formats, three sky condition classes, two contrasting canopy types, and a wide range of canopy cover.

## Methods

### Study sites

Two tree populations from the west side of the Chubut Province (Patagonia Argentina) were selected for this study. Namely, a native *Nothofagus pumilio* forest (broadleaf) and a *Pinus ponderosa* plantation (coniferous). The broadleaf forest is placed on the Huemules Norte experimental unit (42º46′47′′ S, 71º27′ 53′′ W), described in Díaz & Lencinas (2018). This almost primeval forest has a high structural complexity with a top canopy reaching 25 m height. Two stands from the pine plantation were selected, both were pruned and thinned, differing in its average height: 20 m (42°53’9.23”S, 71°20’28.28”W) and 15 m (42º53′49.4′′ S, 71º21′36.5′′ W). Fifteen locations inside each tree population were in-situ selected to ensure a wide range of canopy cover, and they were flagged to ensure repeated acquisition of photographs from the same location (hereafter referred to as photosites).

### Image acquisition and preprocessing

Upward looking photographs were taken with the optical axis vertically aligned and at approximately 1.3 meters from the floor, with shutter delay, and using a tripod. Device characteristics are presented in Fig. 1.

**Fig. 1.**
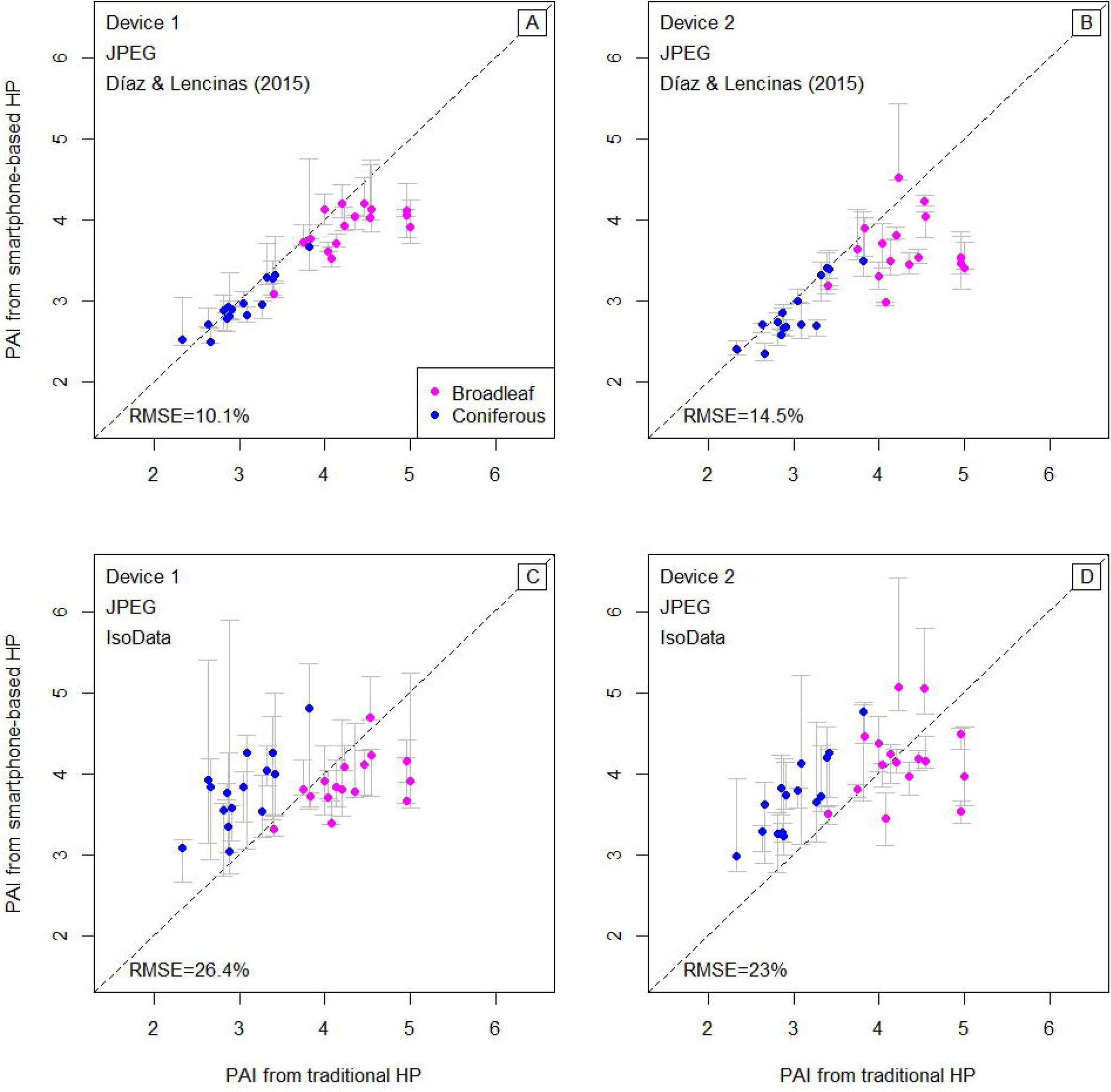
Hemispherical photographs of a *Nothofagus pumilio* forest obtained with the following devises: (A) olloclip auxiliary lens attached to an iPhone 6 Plus—device 1—, (B) Olloclip Multi-Device auxiliary lens attached to a Redmi Note 9—device 2—, (C) and Nikon FC-E9 fisheye converter attached to a Nikon Coolpix 5700 camera—device 3. The field of view (FOV) is annotated on the images, and the lens projection is presented on D.

The Adobe Lightroom App (v.7.1.0 IOS and v.7.2.1 Android) was installed on the smartphones to gain RAW format acquisition and precise exposure manipulation. App features varied with the operating system (OS); under iPhone OS (iOS), exposure compensation can vary from -3 to 3 stop units, while under Redmi-Note-9 OS (Android), from -5 to 5. In the lapse of a few seconds and without moving the smartphone, multiple acquisitions were done with steps of one stop unit, from -3 or -5, depending on OS, and till one stop unit. Smartphones were operated mostly in non-diffuse light conditions (Table 1), so a sunscreen—four-cm-diameter opaque disk attached to a stick and holded by a tripod—was used to avoid direct sunlight reaching the lens. Using Lightroom, the RAW files were exported to sRGB files of 2048 ×1534 pixels in two variants, 8-bit JPEG (quality of 100 %) and 16-bit TIFF (ZIP compression). Since Olloclip lens projection is the same but horizontal FOV varies with smartphone model (Fig. 1), the resolution, expressed as the radius for 90º *θ*, also varies with smartphone model, being 1066 and 910 pixels for device 1 and 2, respectively.

**Table 1.** Acquisition conditions for smartphone-based hemispherical photographs.

| Site | Date (YYYY-MM-DD) | Local time of day (24-hour) | Cloudiness | Wind intensity | Sky condition class |
| --- | --- | --- | --- | --- | --- |
| **La-Zeta** | 2021-12-26 | 13:54 to 16:44 | clear | moderate | Clear |
|  | 2021-12-27 and 30 | 10:58 to 11:52 and 10:58 to 11:37 | cumulus, broken clouds to near complete overcast | moderate | Cloudy |
|  | 2022-03-31 | 15:50 to 17:16 | cirrus, mostly complete bright overcast | light | Overcast |
| **Huemules** | 2022-03-11 | 12:13 to 15:56 | clear | light | Clear |
|  | 2021-12-28 | 14:36 to 17:55 | cirrus, few clouds | light | Cloudy |
|  | 2022-03-31 | 11:53 to 14:19 | cirrus, mostly complete overcast | light | Overcast |

Device 3 was used to acquire in RAW format and in twilight conditions (diffuse light) with the exposure setting manipulated to maximize sensor linear response (Lang et al., 2010). At the La-Zeta site, acquisitions under a clear sky were carried out on 2021-12-23 from 05:54 to 06:27 AM. At the Huemules site, acquisitions were carried out on 2021-12-28 from 8:11 to 9:04 PM, and on 2022-03-11 from 7:23 to 8:37 PM, under a cloudy sky and clear sky, respectively.

Following Lang et al. (2010) methodology, device-3 photographs were developed with the open-source software *dcraw* to avoid gamma correction. However, since device-3 resolution is low (5 megapixels), demosaicing was enabled in *dcraw* instead of selecting the authentic blue spectrum signal from totally RAW data. This was done providing options *-4 -T* to *dcraw*. The result was non-gamma-corrected 16-bit TIFF files of 745 pixels at 90º *θ*. The vignetting effect was corrected (Supplement material S1).

### Image binarization

Photographs acquired with device 1 and 2 were binarized with the Díaz & Lencinas (2015) and the Ridler & Calvard (1978) methods, the former implemented in the *ootb_obia()* function of the *rcaiman* package, and the latter in the *auto_thresh* function of the *autothresholdr* package (Landini et al., 2016).

The Ridler & Calvard (1978) method, hereafter referred to as IsoData, works as a baseline. Jonckheere et al. (2005) identified it as the most robust algorithm for binarizing hemispherical photographs and, since then, it was used as a baseline in many studies, as Cescatti (2007), Loffredo et al. (2016), and Díaz & Lencinas (2018). Following standard practices, the back-gamma corrected blue channel was used (Cescatti, 2007; Díaz & Lencinas, 2018; Lang et al., 2010).

The Díaz & Lencinas (2015) method is a complex chain of algorithms that consumes considerable computing time. Briefly, it starts pixel-wise using the CIELAB color space to produce a synthetic layer in which the blue chroma, no matter its lightness, takes a high score, as well as achromatic pixels with a high lightness; then, detects the sunlit canopy by recognizing the warm greens; and, finally, binarizes the synthetic layer by computing a threshold with IsoData and the synthetic values that do not belong to the sunlit canopy. Next, the processing chain continues segment-wise to detect plant pixels misclassified as sky along the sky-canopy border. The key to this step is to take advantage of the ratio of the blue pixel values to its maximum value at the segment level, which allows handling sky heterogeneity.

Preprocessed images from device 3 were binarized with a revised version of the algorithm presented in Díaz & Lencinas (2018) (Supplement material S1 shows the validation of this method with semi-direct data). Results from device-3 data were the reference values; therefore, the comparison of SHP was against a validated processing chain.

### Retrieval of canopy parameters

A sky grid with cells of 15º of *θ* was used to extract GF from the binarized images. The root mean square error (RMSE) was chosen to summarize the gap fraction errors at the cell-level.

Canopy openness (CO) was calculated with Equation 1 (Gonsamo et al., 2011).

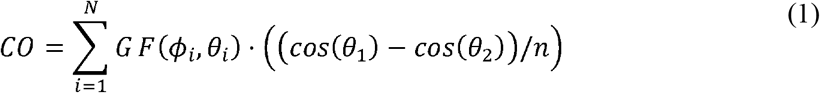

where *GF*(*ϕ*_*I*_,*θ*_*i*_)is the gap fraction of the cell *i, θ*1 and *θ*2 are the smallest and largest zenith angle of the cell *i*, and *n* is the number of cells on the ring delimited by *θ*1 and *θ*2 —zenith range from 20º to 70º. The CIMES package (Gonsamo et al., 2011) was used to calculate ePAI (Miller, 1967) and *Ω* (Chen & Cihlar, 1995). To that end, images from all devices were reprojected to the equidistant projection with a size of 1000 × 1000 pixels, and the same sky region was evaluated across devices.

## Results and discussion

Results from JPEG and TIFF files were almost identical, indicating that color depth does not impact accuracy. Nevertheless, RAW acquisition is recommended since flexibilizes field exposure determination (Macfarlane et al., 2014). Results from TIFF files were made available on Supplementary material S2 only.

Due to its practicality, the histogram shape criteria was selected as the guiding principle for optimal exposure determination. To support the development of a protocol suited for SHP based on that principle, the photograph with the least RMSE per photosite and sky condition class was selected. As a result, four groups were formed (Fig. 2; Supplementary material S3). For each group, the median histogram of the blue channel was computed, and the chi-square distance between the median histogram and each photograph’s histogram was calculated.

**Fig. 2.**
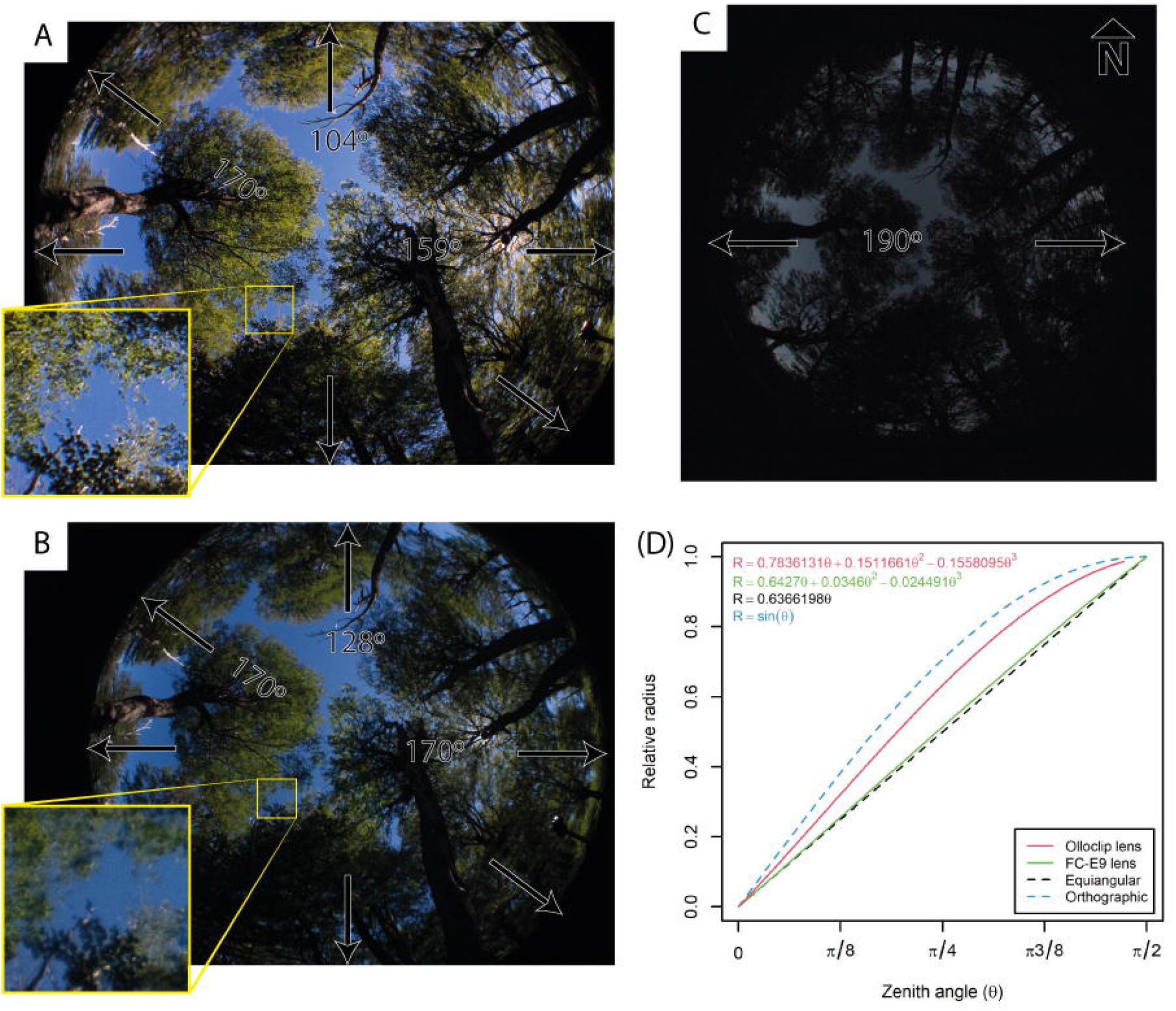
Error distribution of the group formed by the best photograph per photosite and sky condition class. RMSE stands for root mean squared error.

Afterward, the photograph with the least distance per photosite and sky condition class was chosen from the dataset. This is essentially what can be done in the field but with a benefit of objectivity. In the field, the histogram can be inspected to pursue one with wide use of the dynamic range, but very low density of saturated pixels (Fig. 3), as recommended for traditional HP (Macfarlane et al., 2014). As Fig. 4 shows, the median histogram shapes slightly vary with binarization methods and smartphone models, but the mentioned recommendation holds true, although only approximately for device 1 and IsoData.

**Fig. 3.**
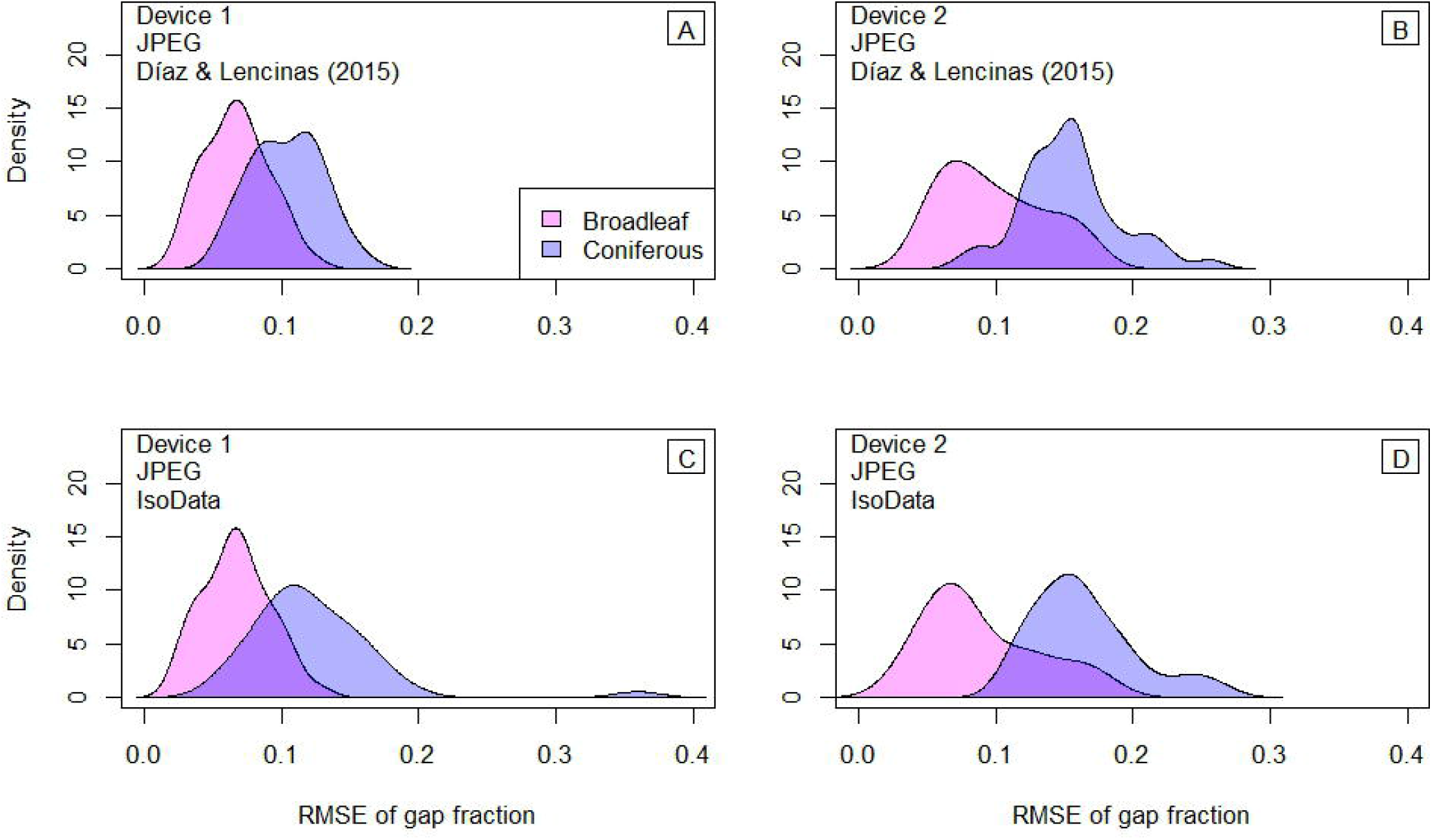
Histogram inspection in Adobe Lightroom App. A–C shows an open canopy, while D–F, a dense canopy. A&D shows overexposed skies; B&E, adequately exposed skies; and C&F, underexposed skies.

**Fig. 4.**
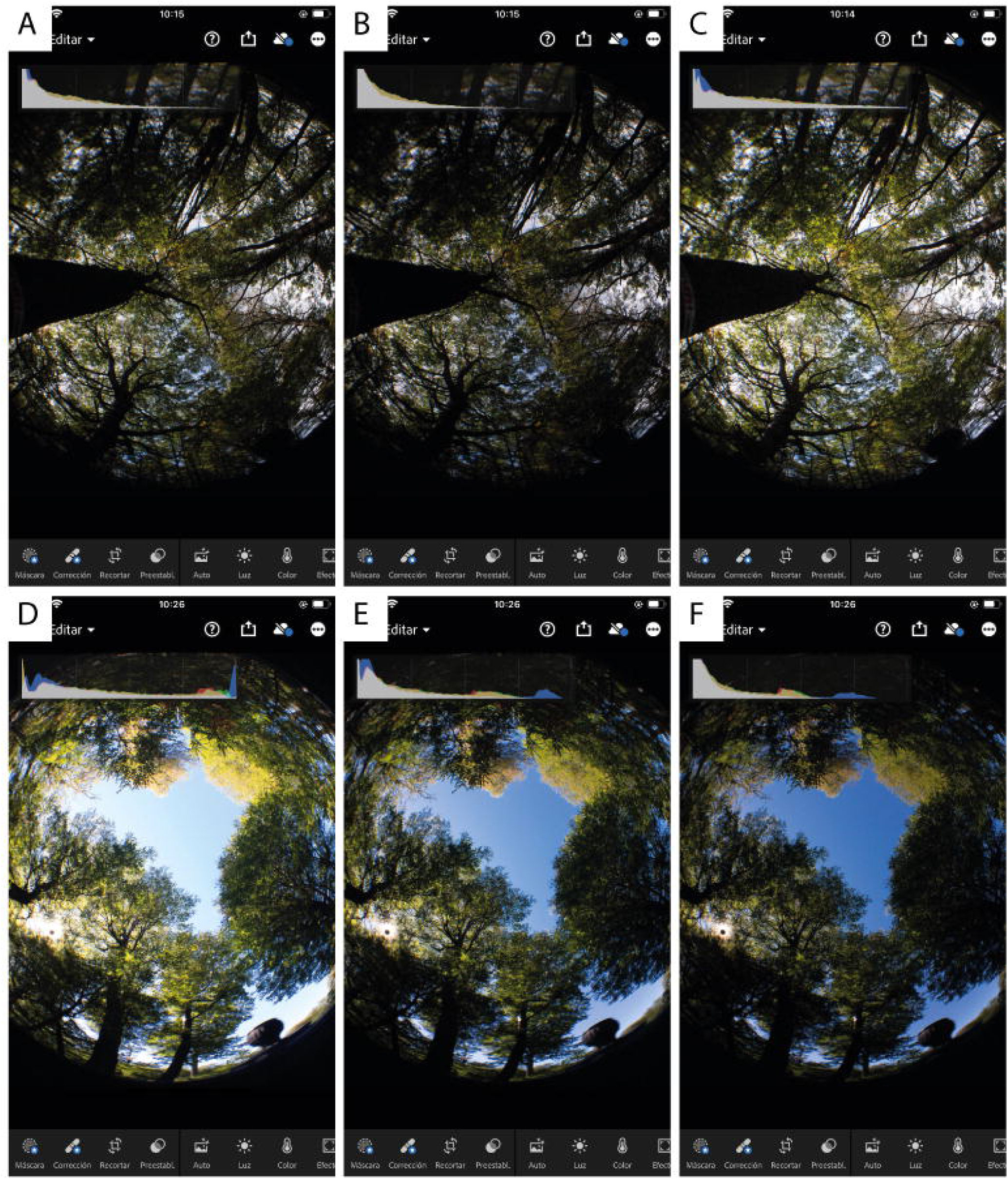
Reference histograms built from the group formed by the best photograph per photosite and sky condition class. Solid lines are the median, and dashed lines are first and third quartiles.

Fig. 5 shows CO calculated from photographs selected with the chi-squared distance criteria. For the coniferous plantation, Díaz & Lencinas (2015) method delivered the most reliable results for all sky conditions. Device 2 did not perform well for the broadleaf forest, independently of the binarization method. For device 1 and broadleaf forest, both methods delivered good results.

**Fig. 5.**
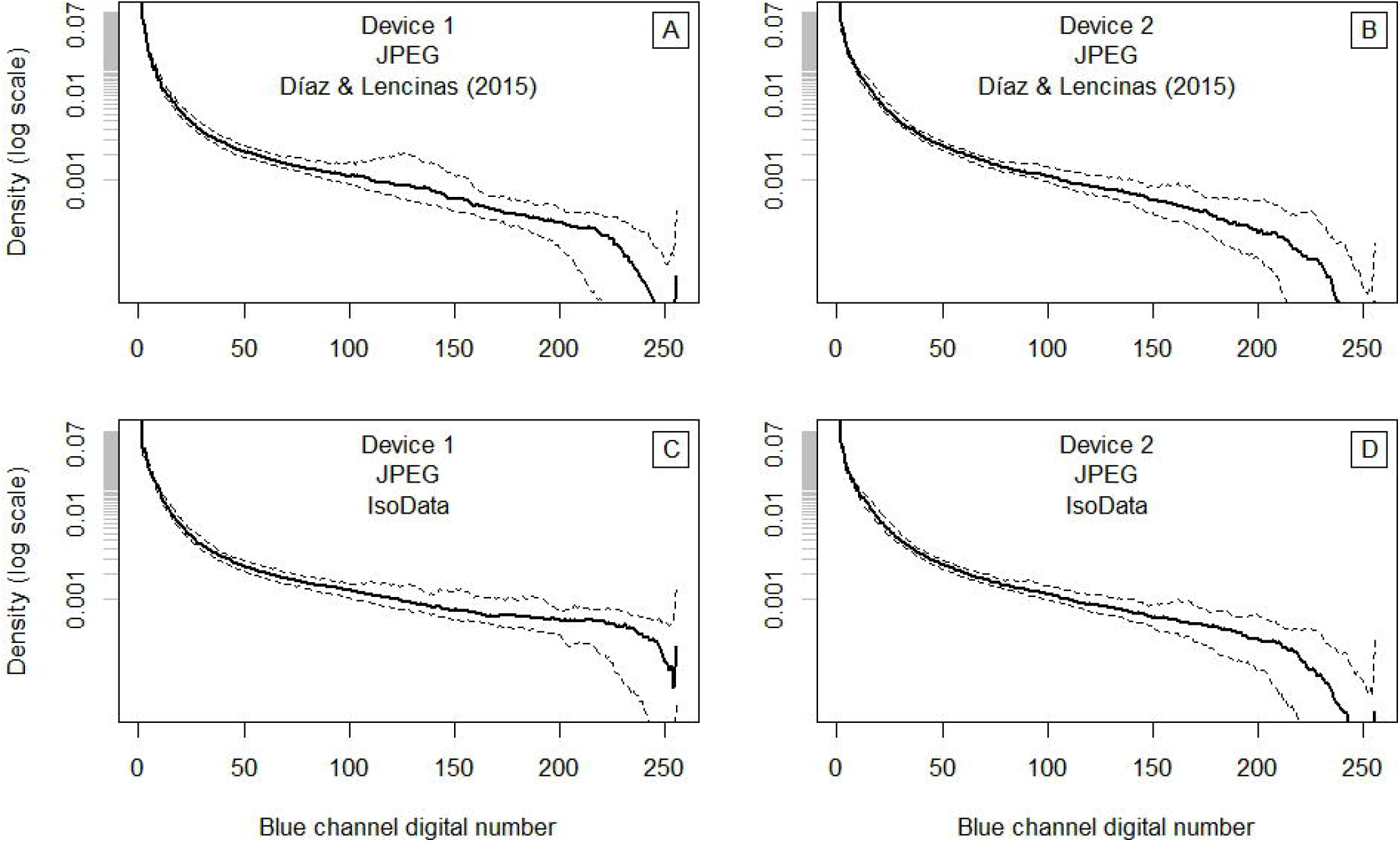
Accuracy assessment of smartphone-based hemispherical photography through canopy openness (CO) comparison. RMSE stands for root mean square error.

Similar to Fig. 5, Fig. 6 shows PAI calculated with photographs selected with the chi-squared distance criteria, but summarizing instead of discriminating between sky condition classes— the RMSE shown in the figure was calculated before summarizing. The better sharpness of device 1 relative to device 2 (Fig. 1) seems to have an impact on PAI accuracy. However, the most notorious difference was between binarization methods, suggesting that Díaz & Lencinas (2015) is indeed more accurate than IsoData (RMSE of 12% vs. 25%) and can produce good results (RMSE ~ 10%) from well-exposed and sharp images, independently of light conditions and forest type. It is worth noting that a trend of overestimation in the extremely open canopies and underestimation in the extremely closed canopies is observed in Fig. 6A. This seems a consequence of using the shape of the median histogram as reference for the chi-distance criteria because histogram shape changes substantially with CO (Fig. 3). Moreover, errors in the extremes are lower when the RMSE criteria is used to select the best photograph per photosite or when histogram is manipulated before RAW development (Supplementary materials S4).

**Fig. 6.**
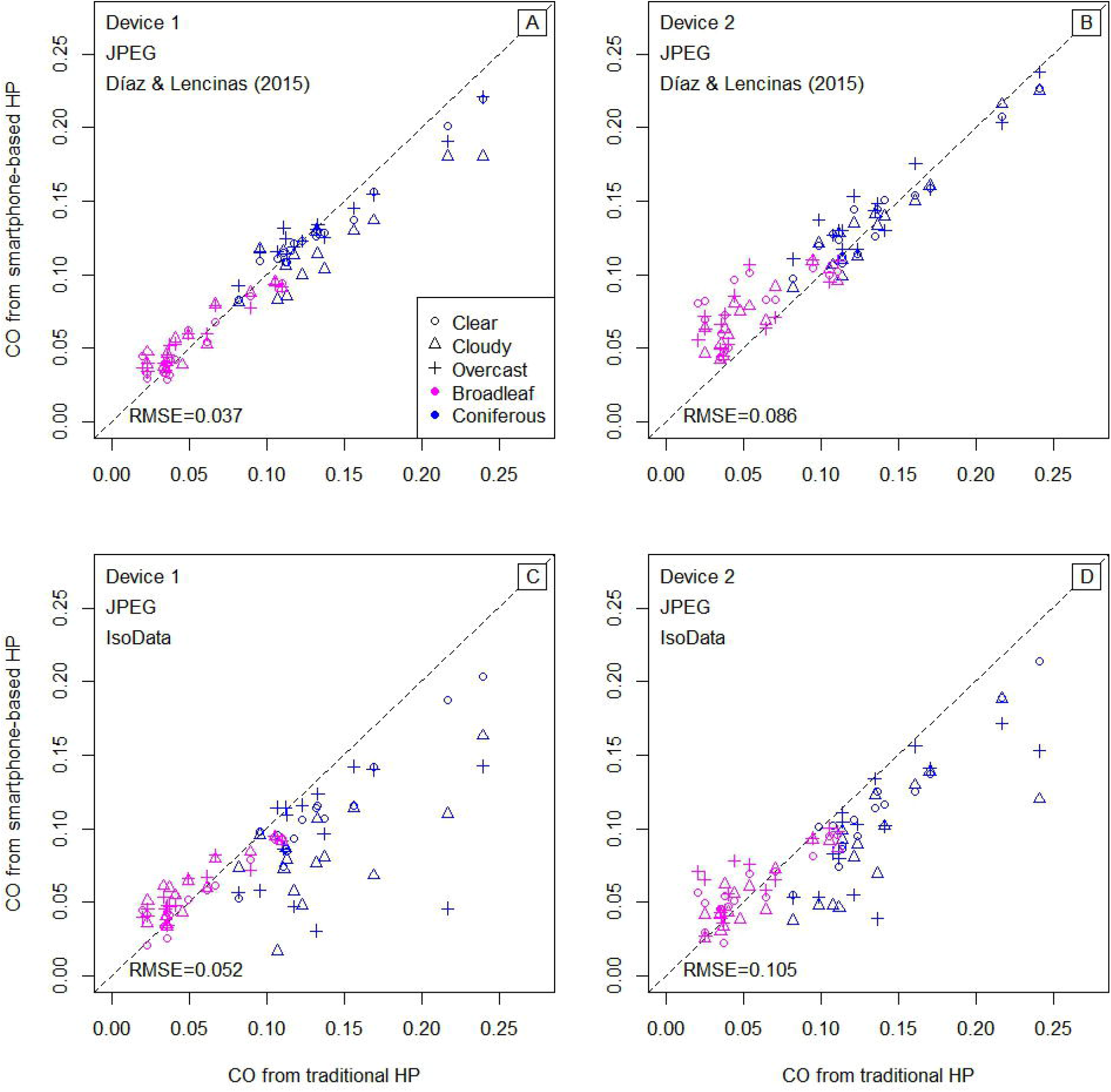
Accuracy assessment of smartphone-based hemispherical photography through plant area index comparison. Points are the median of three values (different sky conditions) and whisker indicates the range. RMSE stands for root mean square error.

Results indicate that image fidelity can be characterized by sharpness (determined by the device) and sky color fidelity (controlled through exposure manipulation). Therefore, if portability is not an issue, professional-grade equipment can be used to take advantage of better image fidelity. Regarding other procedure details, refer to Cameron et al. (2021).

## Conclusion

Canopy hemispherical photography assumes lens projection is known, image fidelity is high, and binarization is accurate. Therefore, a proper lens calibration is mandatory to obtain the degree of accuracy reported here, as well as sharpness and color fidelity. Having means to precisely manipulate the exposure and evaluate histogram shape is required to ensure wide use of the dynamic range but very low density of saturated pixels (the Adobe Lightroom app is a viable alternative). This will be translated to an adequate sky color fidelity, which could be quickly evaluated on the field by using basic image interpretation skills. Finally, if an image obtained with the aforementioned precautions is provided to the binarization method by Díaz & Lencinas (2015), it will likely provide reliable results independently of sky conditions.

## Data accessibility

The ‘rcaiman’ package is available on CRAN repository. Data are available in an OSF repository (a-link-will-be-provided-after-acceptance).

