## Supplementary material for "Optimizing forest canopy structure retrieval from smartphone-based hemispherical photography": Suplementary materials

The present supplementary material is related to the article entitled "Optimizing forest canopy structure retrieval from smartphone-based hemispherical photography," by Gastón Mauro Díaz.

#### S1. Reprocessed data from Díaz & Lencinas (2018)

Díaz & Lencinas (2018) presents a new method, hereafter called model-based local thresholding (MBLT), and its validation against semidirect leaf area index (LAI) calculated from litter-traps data.

In essence, the MBLT approach proposes a linear relationship between background value and optimal threshold value. The current implementation uses a statistical-driven reconstruction of the canopy photograph background (i.e., the sky). The statistical models used for sky reconstruction are able to explain smooth changes in sky brightness, so the approach works best under clear skies or overcast conditions. After the reconstruction, the local threshold is linearly predicted from sky brightness.

The data used in Díaz & Lencinas (2018) is from ten plots of 10 m radius carefully selected to cover the structural variability found in the Huemules site (refer to main text). In each plot, three 65-cm-diameter litter traps were randomly located and four photographs taken, one in the center and three over the plot perimeter and azimuth 0º, 120º, and 240º.

Device 3, described in the main document, was employed to take the photographs. The chosen file format was JPEG, and the exposure was manipulated to produce a series of photographs per photosite. In 2021, the vignetting effect from device 3 was measured with a photometric sphere at Tartu Observatory, and a vignetting correction function was developed, which was applied to the JPEG files as a preprocessing step.

All photographs were binarized with the *ootb_mblt* from the *rcaiman* package (refer to main text). Afterward, using the respective binarized image as a sky mask, the median sky brightness (blue channel, gamma-back corrected) was obtained and used to select the photograph per photosite with sky brightness closer to 0.5. The latter allowed the selection of the binarized images from photographs with a constant exposure relative to the open sky auto-exposure (Zhang et al., 2005) since exposure controls brightness (Macfarlane et al., 2014; Díaz & Lencinas, 2018).

The retrieval of plant area index (PAI) from the binarized images was done as described in the main document. However, instead of using the four photosites per plot and the whole azimuthal range per image, only photosites from the perimeter were selected, and an azimuthal range of 60º oriented to the plot center was used to mask the binarized images.

Finally, the PAI was averaged per plot and compared against the semi-direct leaf area index from litter-trap data (Fig. I).

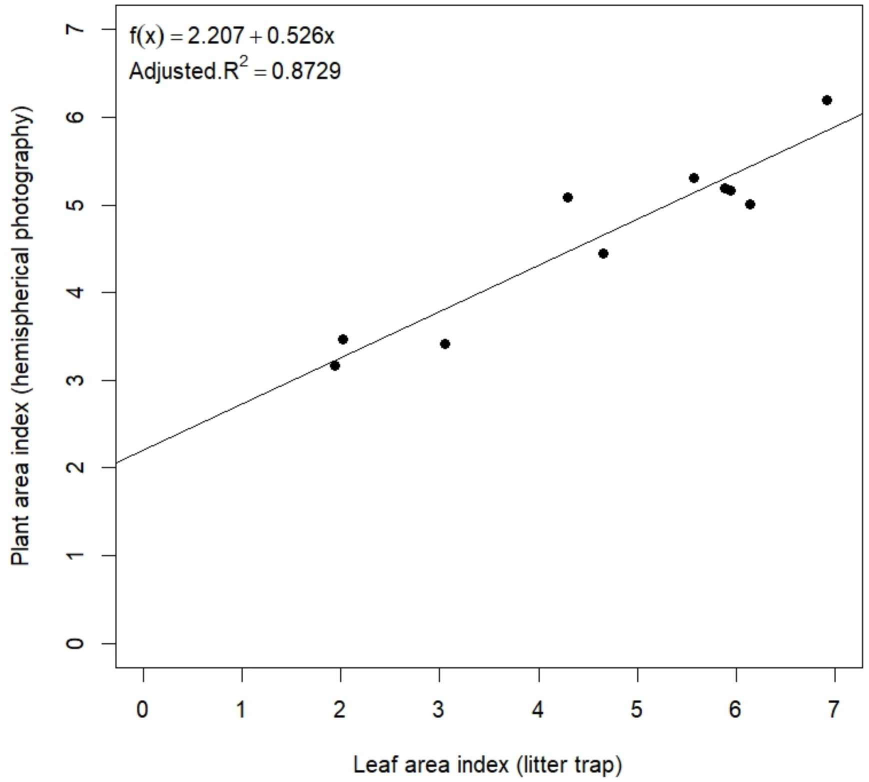


Fig. I. Validation of the method used as reference in the main research. The line is the fitted linear model.

### S2. Figures of TIFF-files data

Since results from JPEG and TIFF files were almost identical, the formers were shown in the main document, while the latters here. This choice was driven by practicality since JPEG is broadly used and has effective data compression. It is worth mentioning that developing the RAW files with *dcraw,* as it was done for device 3 (refer to main document), did not lead to good quality pictures.

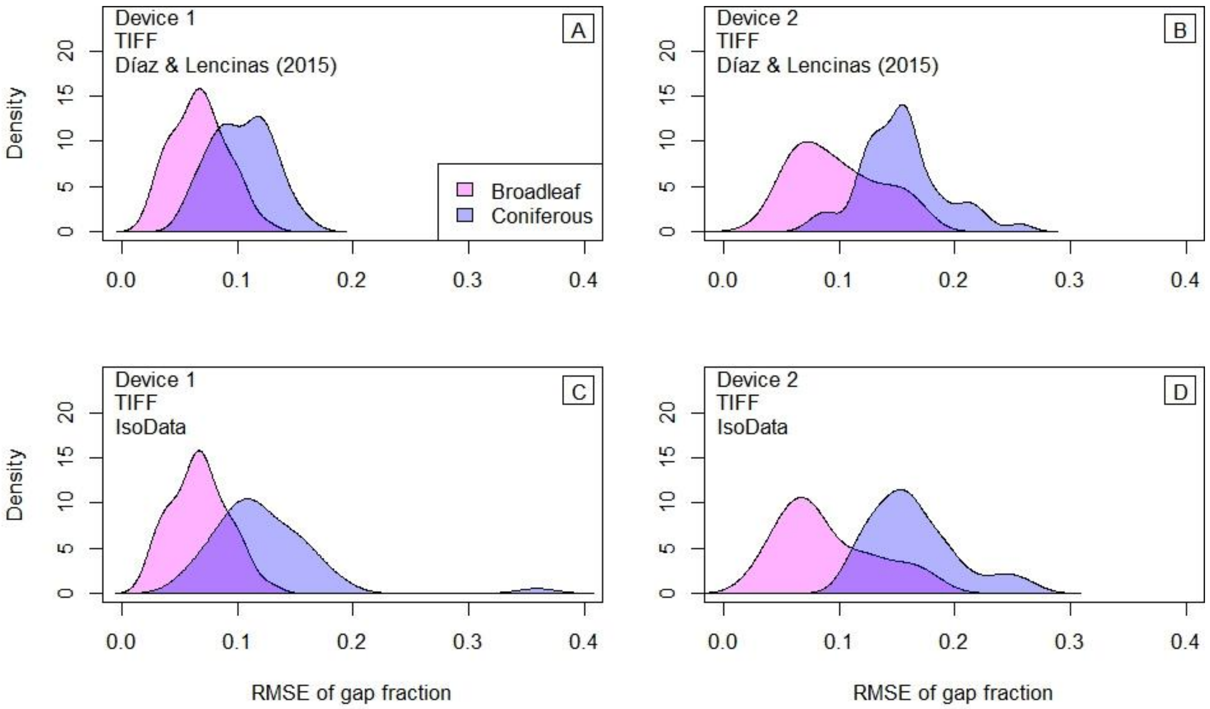


Fig. 2S. Error distribution of the group formed by the best photograph per photosite and sky condition class. RMSE stands for root mean squared error.

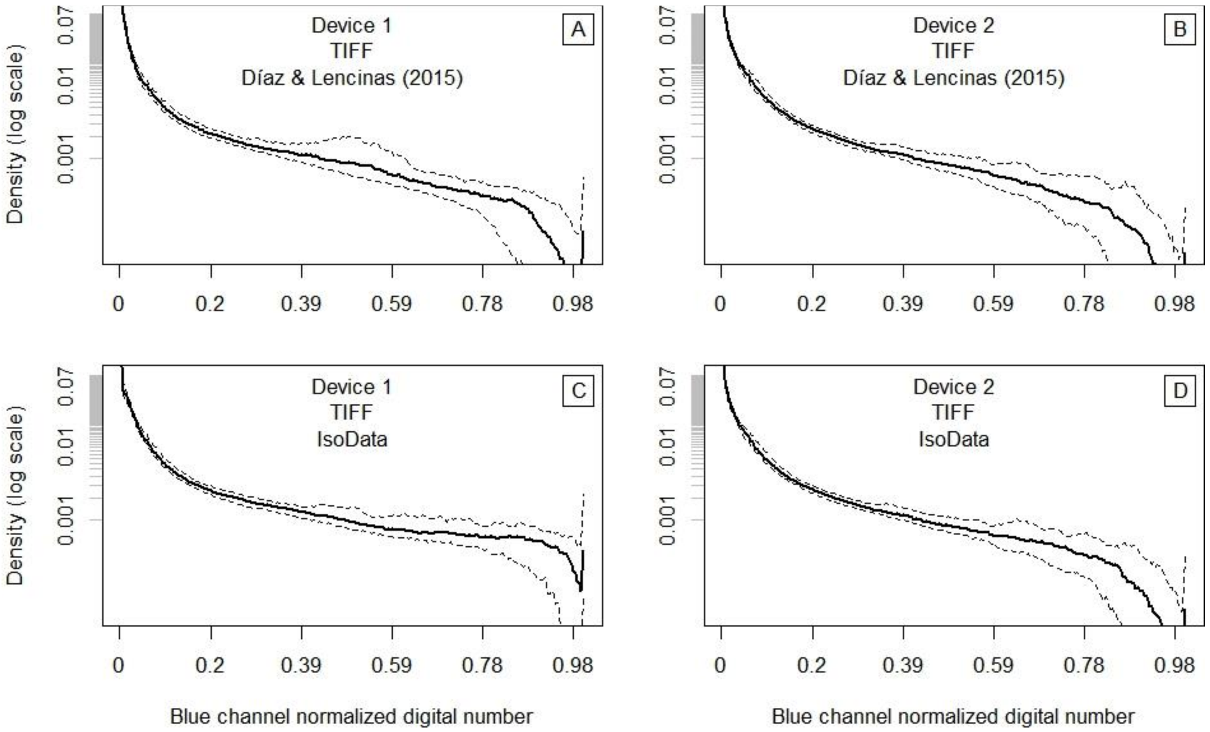


Fig. 4S. Reference histograms built from the group formed by the best photograph per photosite and sky condition class. Solid lines are the median, and dashed lines are first and third quartiles.

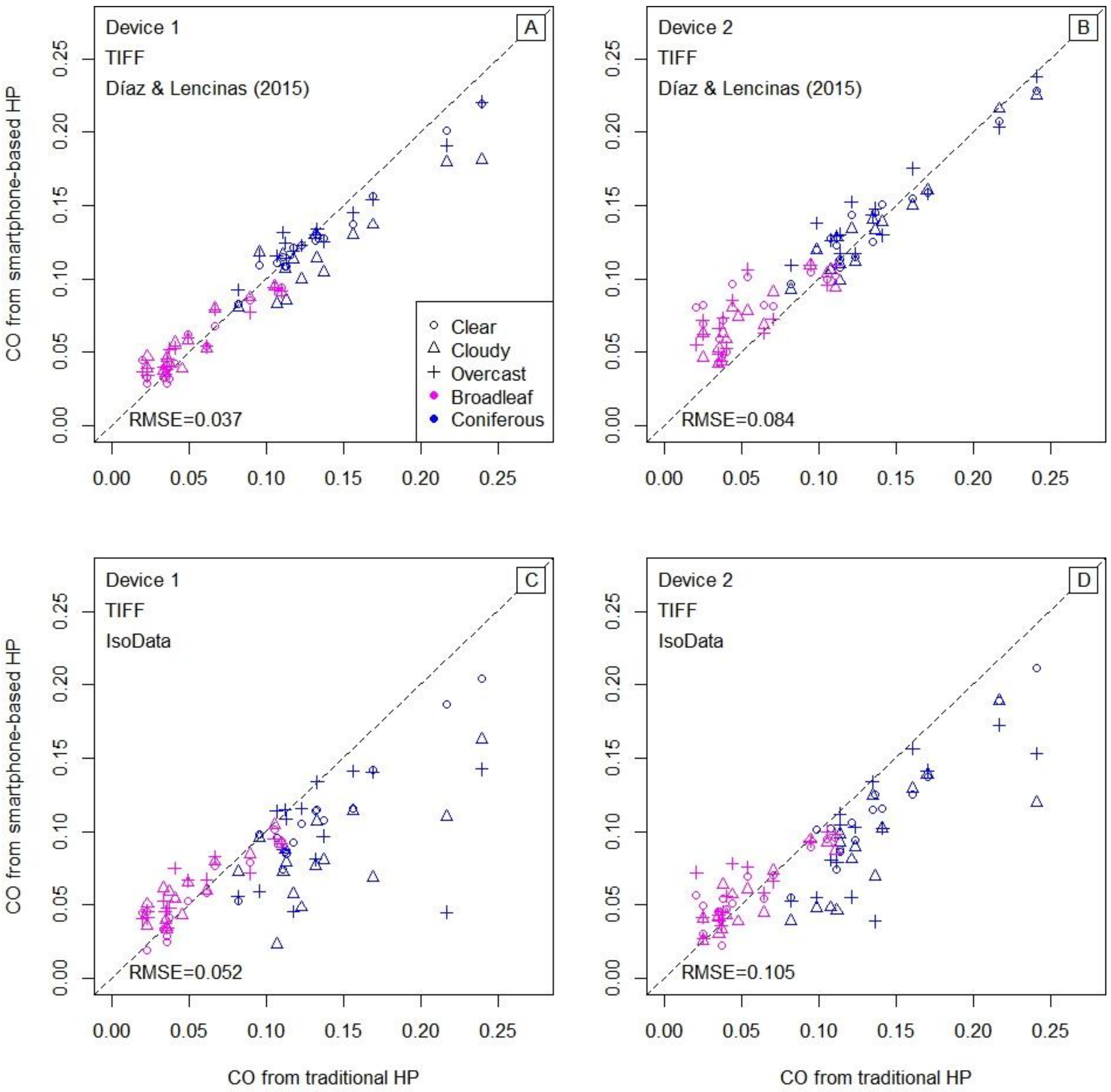


Fig. 5S. Accuracy assessment of smartphone-based hemispherical photography through canopy openness (CO) comparison. RMSE stands for root mean square error.

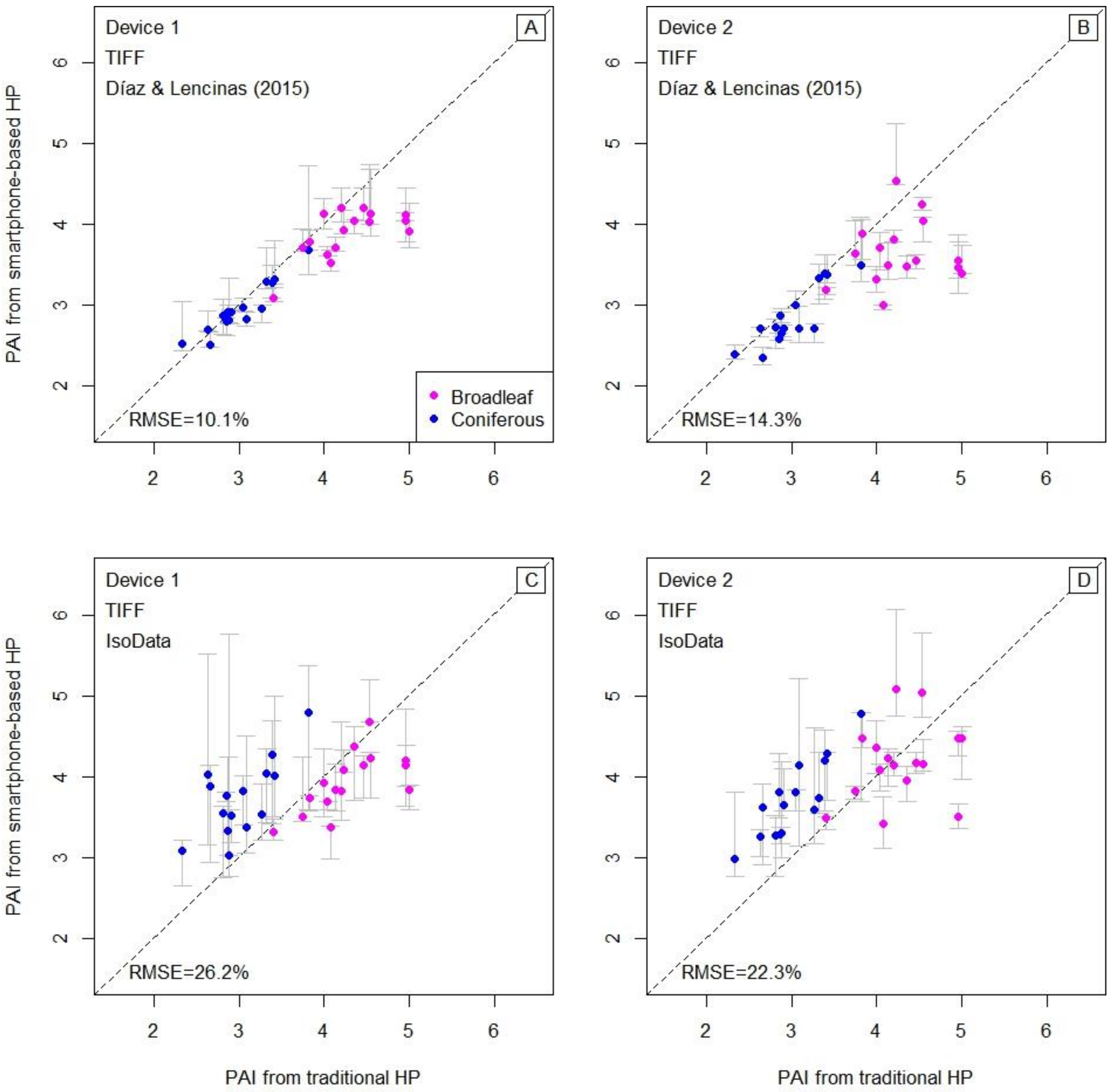


Fig. 6S. Accuracy assessment of smartphone-based hemispherical photography through plant area index comparison. Points are the median of three values (different sky conditions) and whisker indicates the range. RMSE stands for root mean square error.

### S3. Contact-sheet-like images

Contact-sheet-like images of the photographs selected with the RMSE criteria (refer to the main document). Figures II to V have been built as a matrix of 10 columns and 9 rows. Columns 1 to 5 are La-Zeta data, and 6 to 10 are Huemules data. Rows 1 to 3 belong to the class “clear”, 4 to 6 to “cloudy”, and 7 to 9 to “overcast”.

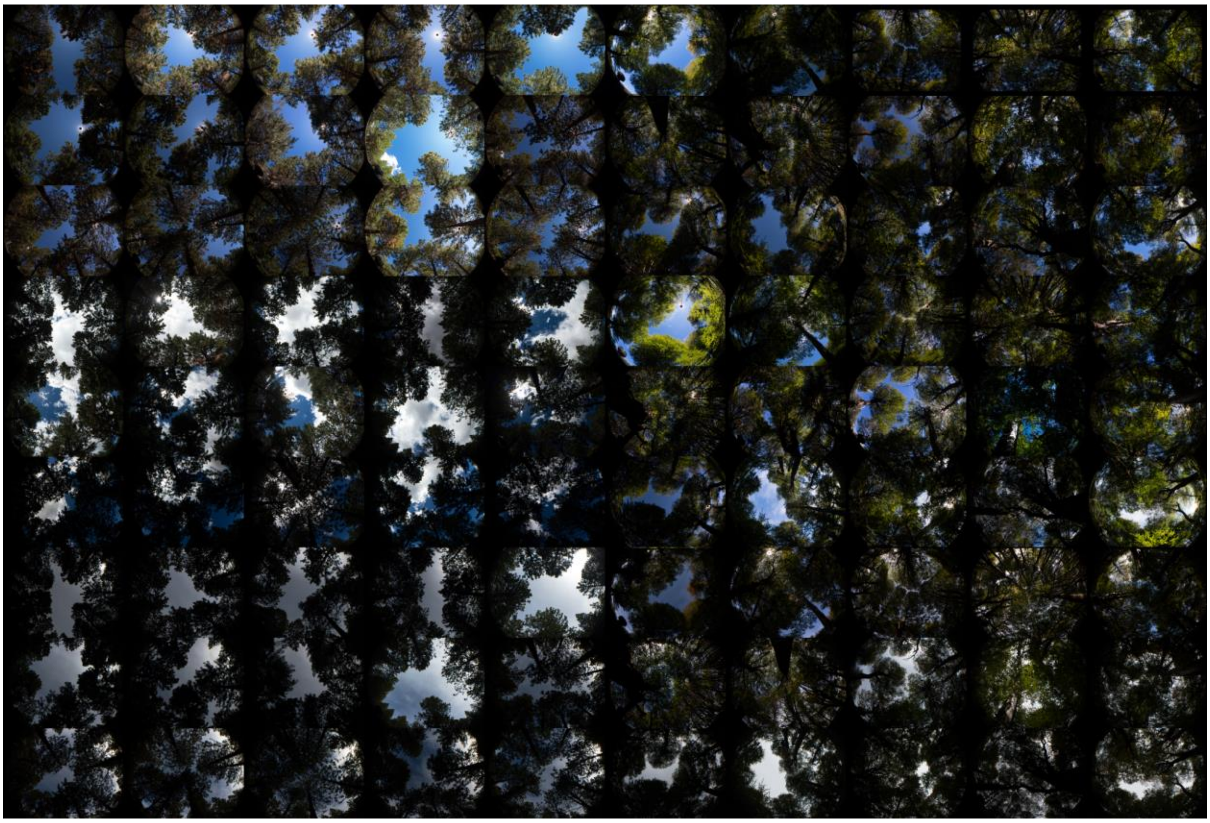

Fig. II. Photographs from device 1 selected with the RMSE after processing them with Díaz & Lencinas (2015) method.

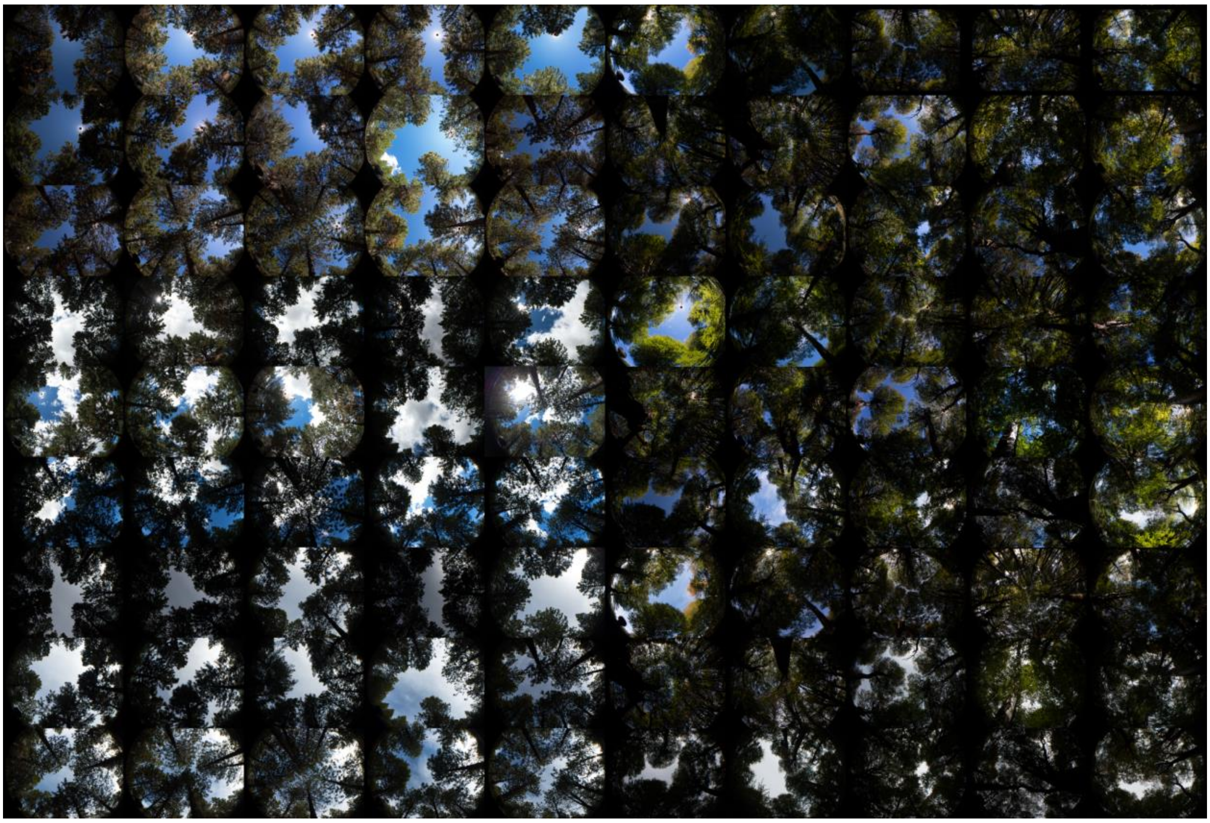

Fig. III. Photographs from device 1 selected with the RMSE after processing them with the IsoData method.

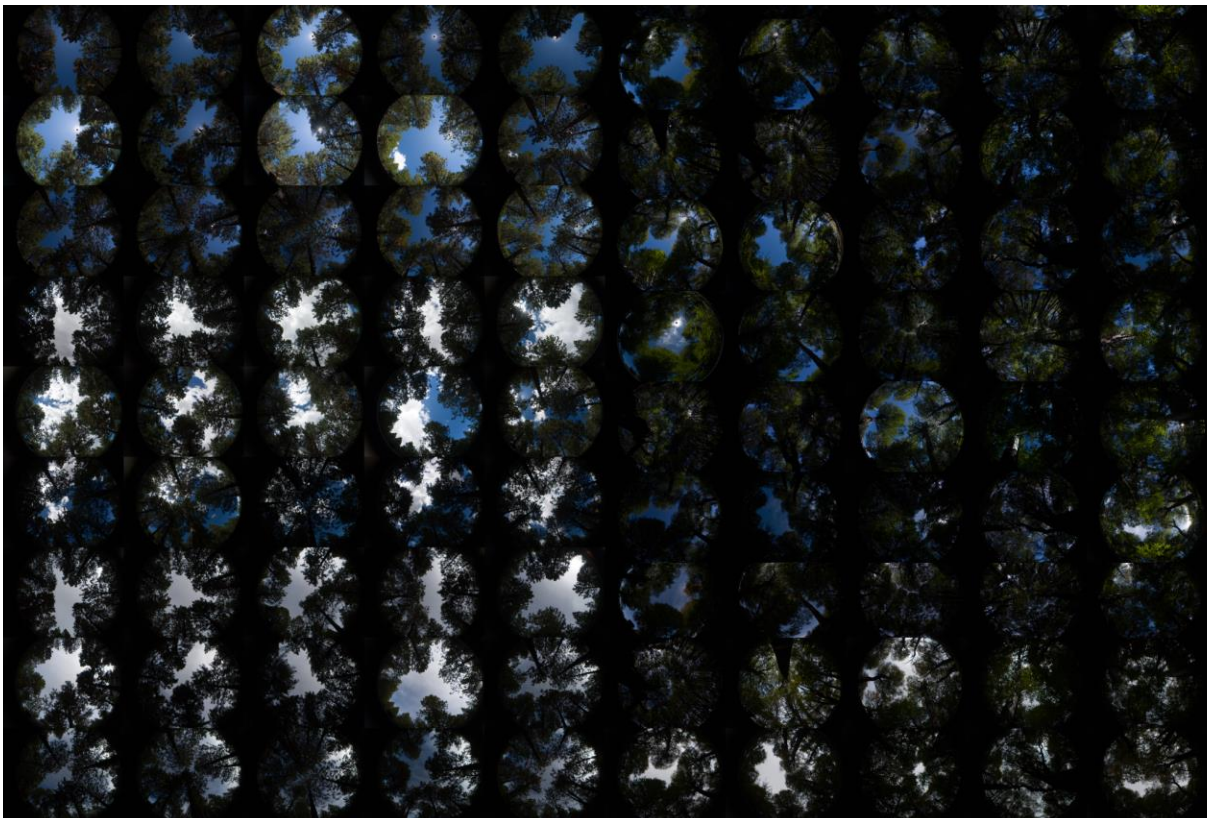

Fig. IV. Photographs from device 2 selected with the RMSE after processing them with Díaz & Lencinas (2015) method.

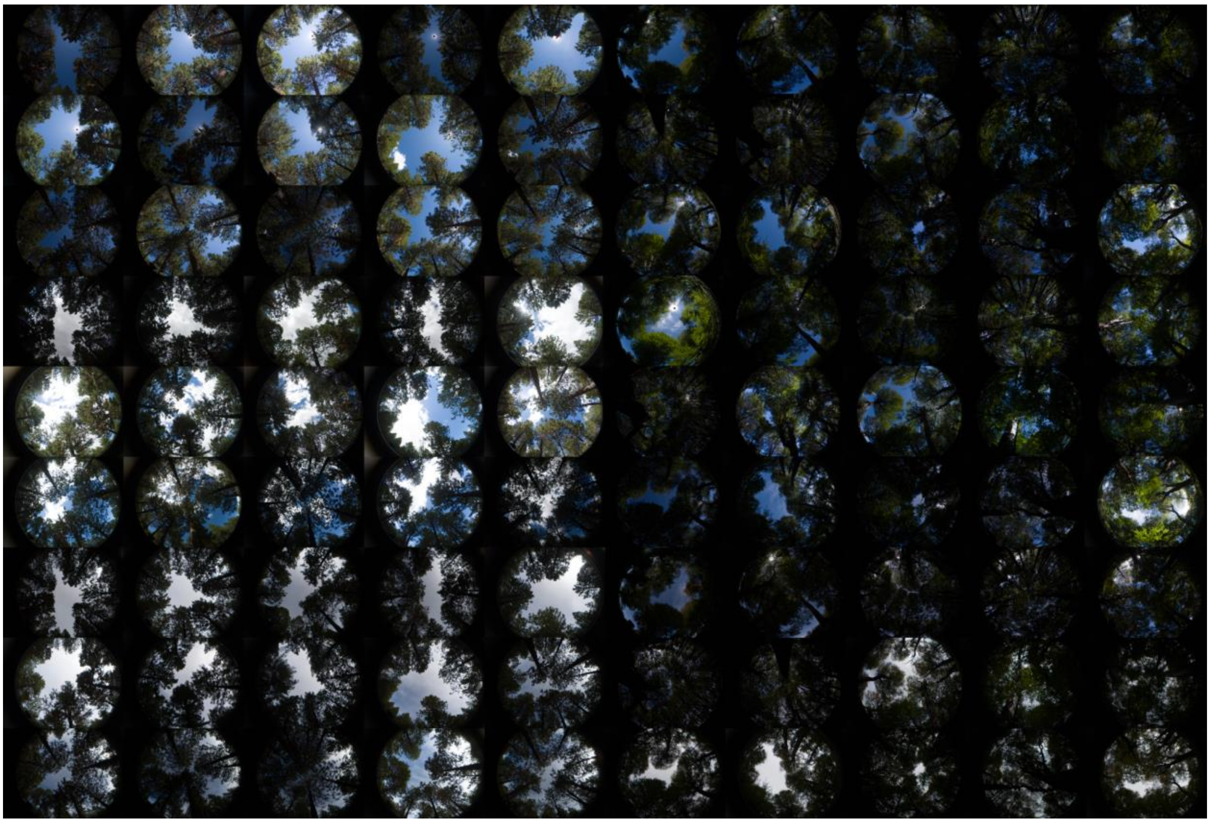

Fig. V. Photographs from device 2 selected with the RMSE after processing them with the IsoData method.

Theoretically, IsoData will work best with homogeneous backgrounds because it is a single threshold applied to the whole picture. To some extent, this can be seen through a visual comparison of Fig. II against Fig. III, and Fig. IV against Fig. V., particularly La-Zeta data and "overcast".

### S4. Potential accuracy

To produce Fig. VI, photos were manually selected from the dataset after completing the research and drawing conclusions, replicating what an informed technician could accomplish. Therefore, Fig. VI shows the accuracy that a technician can obtain by acquiring photographs in RAW format with the recommended protocol and adjusting their histogram, if needed, before developing them to JPEG format.

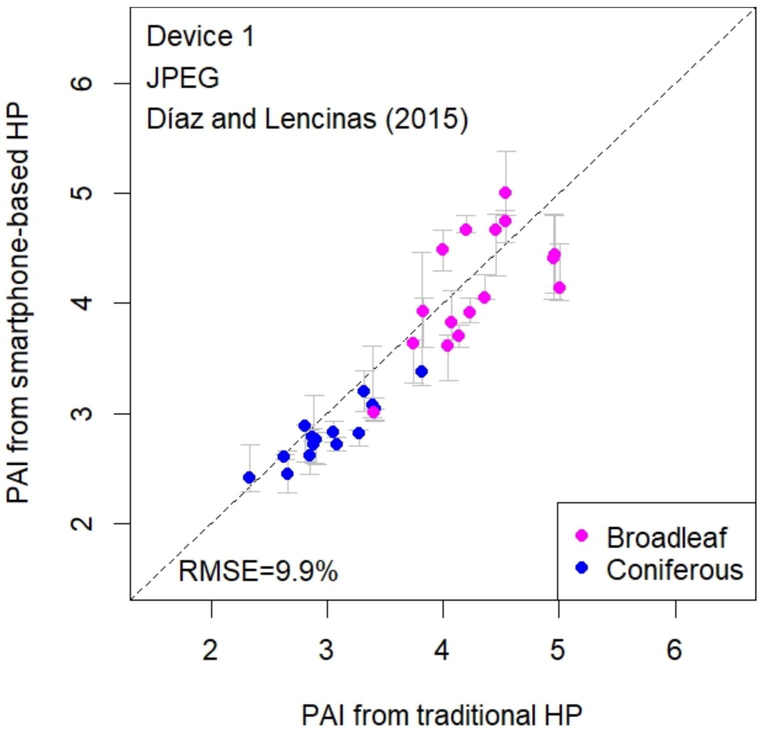


Fig. VI. PAI calculated after manually selecting, for each photosite and sky condition class, the JPEG file with the highest image fidelity (according to the main document recommendations). Points are the median of three values (different sky conditions) and whisker indicates the range. RMSE stands for root mean square error.

Fig. VII shows the PAI calculated with the photographs on the dataset having the most accurate gap fraction. This potential accuracy shows that the device can record the canopy very well, although deliberately producing the required level of image fidelity is challenging, as the experience described in the previous paragraph demonstrates.

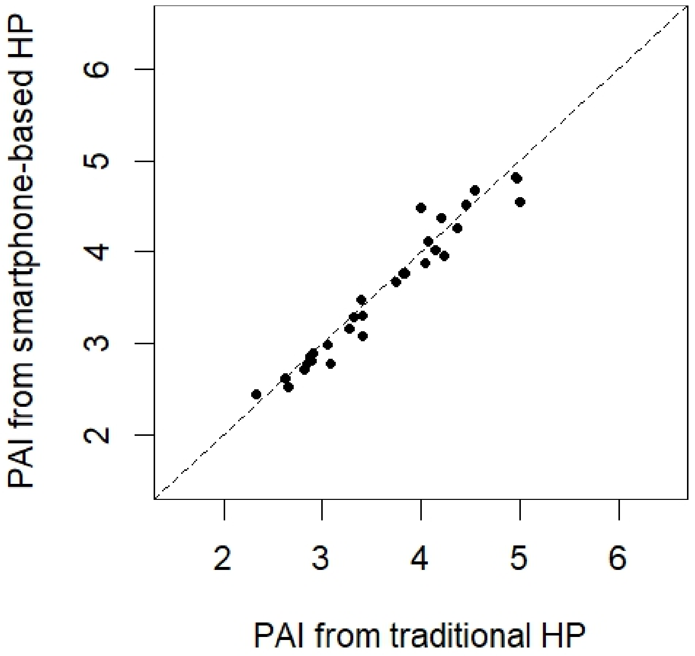


Fig. VII. PAI calculated after selecting, for each photosite, the JPEG file from device 1 that was binarized with the Díaz & Lencinas (2015) method and has the gap fraction closer to the reference data. Please note that this level of accuracy may be impossible to reach outside experimental conditions when there is no reference data to be used as criteria for choosing the best photograph.
